# Generalised Complementarity Analysis: identifying the most precious places for the conservation of Species, Functional and Phylogenetic Diversity

**DOI:** 10.1101/189837

**Authors:** David Anthony Nipperess

## Abstract

The most precious places for conservation are those that make the largest contribution to regional, national or global biodiversity. The two key concepts for determining the contribution of a specific site are *Complementarity* (the gain in diversity achieved when adding that site to a set of other sites) and *Irreplaceability* (here defined as the overall complementarity of that site when compared to a range of possible combinations of other sites). *Generalised Complementarity Analysis* (GCA) is a mathematical framework that provides an exact analytical solution for the expected complementarity (gain in diversity) of a focal site, when added to a set of other sites of a given size (*m*). Diversity is defined very generally to allow for complementarity to be calculated for species richness, Functional Diversity or Phylogenetic Diversity. The expected irreplaceability of a focal site is then defined in GCA as the area under the curve of expected complementarity values for all possible values of *m*. GCA is much more computationally efficient than existing algorithmic approaches and is scalable to very large numbers of sites. Because complementarity and irreplaceability are calculated for all possible combinations of sites, GCA serves as a null model for systematic conservation planning algorithms that seek to optimise site selection. However, because truly irreplaceable sites remain so under all possible site selections, GCA is a powerful conservation planning tool in its own right, providing an efficient means of identifying the world’s most precious places for conservation.

## Introduction

If we are to conserve biodiversity, we must do so efficiently while still ensuring that all species (or other units of biodiversity) are adequately protected. When selecting sites for protection, conserving biodiversity becomes a problem of representativeness: does some set of sites adequately represent biodiversity regionally, nationally or globally? This general problem is the impetus for the science of systematic conservation planning, which aims to find optimal solutions to site selection and prioritisation for biodiversity conservation (Margules & Pressey 2000).

*Complementarity* and *Irreplaceability* are two central concepts in systematic conservation planning. Complementarity is defined as the gain in diversity of adding some site to an existing set of sites (Williams et al. 2006). Conservation planning software, such as Marxan (Ball et al. 2009) and Zonation (Moilanen et al. 2005), use complementarity as a principal criterion to search for combinations of sites that optimise the conservation of biodiversity for the least cost (typically, the smallest number of sites). Irreplaceability is then defined as the importance of a site in ensuring that all species (or some other target) are conserved, usually quantified as the frequency with which a site is included in an optimal combination of sites (Pressey et al. 1994).

Because the number of possible combinations of sites is potentially very large, conservation planning algorithms are computationally intensive and must employ heuristic methods that search through the potential options without assessing all possible solutions (combinations of sites). As a result, producing fine-scale maps of conservation importance (i.e. irreplaceability) is still challenging, despite rapid increases in computer power. Further, because not all possible solutions are assessed, it is not feasible to determine the overall importance of a site for biodiversity conservation, across all possible scenarios, even though this would be very useful information and would provide more flexibility in conservation decision making.

Conservation planning also does not commonly take account of species differences despite the wide appreciation that all species do not contribute equally to biodiversity. In a similar way to sites, some species are more irreplaceable than others by having traits (ecological, behavioural, physiological) that are relatively rare. It follows that the irreplaceability of sites is not only a consequence of the geographic rarity of the species present but also the uniqueness of those species themselves. Uniqueness can be measured in terms of species traits or in an evolutionary sense - species that are closely related can be reliably assumed to be more similar for a variety of traits than more distantly related species. Apart from distinctive traits (which may contribute to ecosystem function), the unique evolutionary history represented by a species is itself worthy of conservation (Rosauer & Mooers 2013). Despite the wide recognition of its importance, information on the similarity or difference between species has been implemented only rarely in conservation planning (Asmyhr et al. 2014; Pollock et al. 2015; Pollock et al. 2017) and full mathematical integration into conservation planning algorithms is still lacking.

Building on previous research on the rarefaction of species richness (Hurlbert 1971), phylogenetic extensions to rarefaction (Chao et al. 2015; Nipperess & Matsen 2013; Nipperess 2016; Tsirogiannis et al. 2012) and complementarity analysis (Williams et al. 2006), I present a novel mathematical framework (*Generalised Complementarity Analysis*) for informing conservation planning. This framework: 1) allows for the exact calculation of complementarity and irreplaceability, given all possible scenarios; 2) is computationally efficient and scalable to very large numbers of sites; and 3) is flexible enough to incorporate information on the phylogenetic or functional relationships between species. Generalised Complementarity Analysis is thus “generalised” in two senses: 1) it calculates complementarity and irreplaceability for all possible combinations of sites (rather than optimal solutions only); and 2) can be based on a variety of diversity measures.

## Mathematical framework

A useful working definition of complementarity is the gain in diversity achieved by adding a specific site to a set of sites (Williams et al. 2006). In other words, complementarity is the contribution of a site to overall diversity. I define diversity (*D*) very generally as some measure capturing information of the biological variety present in the set. However, the diversity measure chosen must have the property of concavity, meaning that the addition of additional elements to a set must never decrease the diversity of the set (Lande 1996). Diversity measures that fulfil this criterion include species richness (and some other forms of Species Diversity), Phylogenetic Diversity (Faith 1992) and Functional Diversity (Petchey & Gaston 2002).

Let *N* (of size *n*) be a set of all possible sites under consideration and let *M* be any subset (of size *m*) drawn from *N*. Complementarity is thus a property of a particular site (*i*) and is the gain in diversity (*ΔD*) of adding the site to some subset of sites (*M*) of which *i* is not a member(*i* ∉ *M* ⊂ *N*) (Nipperess 2016; Olszewski 2004) (equation 1).

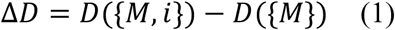

Where *D*({*M*, *i*}) is the total diversity of sites comprising the set, *M*, *and* the site, *i*, and *D*({*M*}) is the diversity of the set, *M*.

If *M* is permitted to be any possible subset of *N* of some given size (*m*), then the expectation of complementarity for any specific site (*i*) can be derived. This is, in effect, a null model for complementarity-based conservation planning algorithms, as it measures mean complementarity for all possible solutions (all combinations of sites of a given size) and not just those that are optimal.

For the simplest case of complementarity in species richness, the expected complementarity (*E*[*ΔD*(*i*,*m*)]) of site *i* is the sum of the probability (*p*_*ij*_) that each species (*j*) present in site *i* is absent from a subset of sites (*M* where *i* ∉ *M*) of size *m*, drawn at random and without replacement from *N* (equation 2).

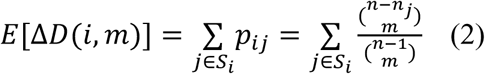

Where *S*_*i*_ is the subset of species present in site *i* (where *S*_*i*_ ⊂ *S* and *S* is the full set of species found in *N*) and where *n*_*j*_ is the frequency with which species *j* occurs across all sites in *N* (i.e. 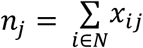 where *x*_*ij*_ is the incidence (absent = 0 / present = 1) of species *j* in site *i*). Note that *n* – *n*_*j*_ = (*n* – 1) – (*n*_*j*_ – 1) where (*n* – 1) is the size of the set containing all sites in *N except* site *i*, and (*n*_*j*_ – 1) is the number of sites in *N* that are *not* site *i* but where species *j* also occurs.

Expected complementarity (equation 2) bears a close similarity to the classic analytical solution for rarefaction (Hurlbert 1971) and, like rarefaction, is derived from the hypergeometric distribution. Equation 2 is, in fact, the exact analytical solution for the complementarity analysis of Williams et al. (2006), who used repeated Monte Carlo sampling to derive an estimate of expected complementarity. With an analytical solution, such computer intensive methods are no longer necessary.

Expected complementarity for species richness can be easily adapted for any concave diversity measure that is tree-based. A tree-based diversity measure defines diversity as the total branch length on a tree connecting a set of species via branch segments to the root of the tree, and where difference between a pair of species is measured as the path length between them (Figure 1). Tree-based diversity is a class of measures (Cardoso et al. 2014) that includes Phylogenetic Diversity (PD) (Faith 1992), Functional Diversity (FD) (Petchey & Gaston 2002) and, as a special case, species richness (Nipperess & Matsen 2013).

**Figure 1:**
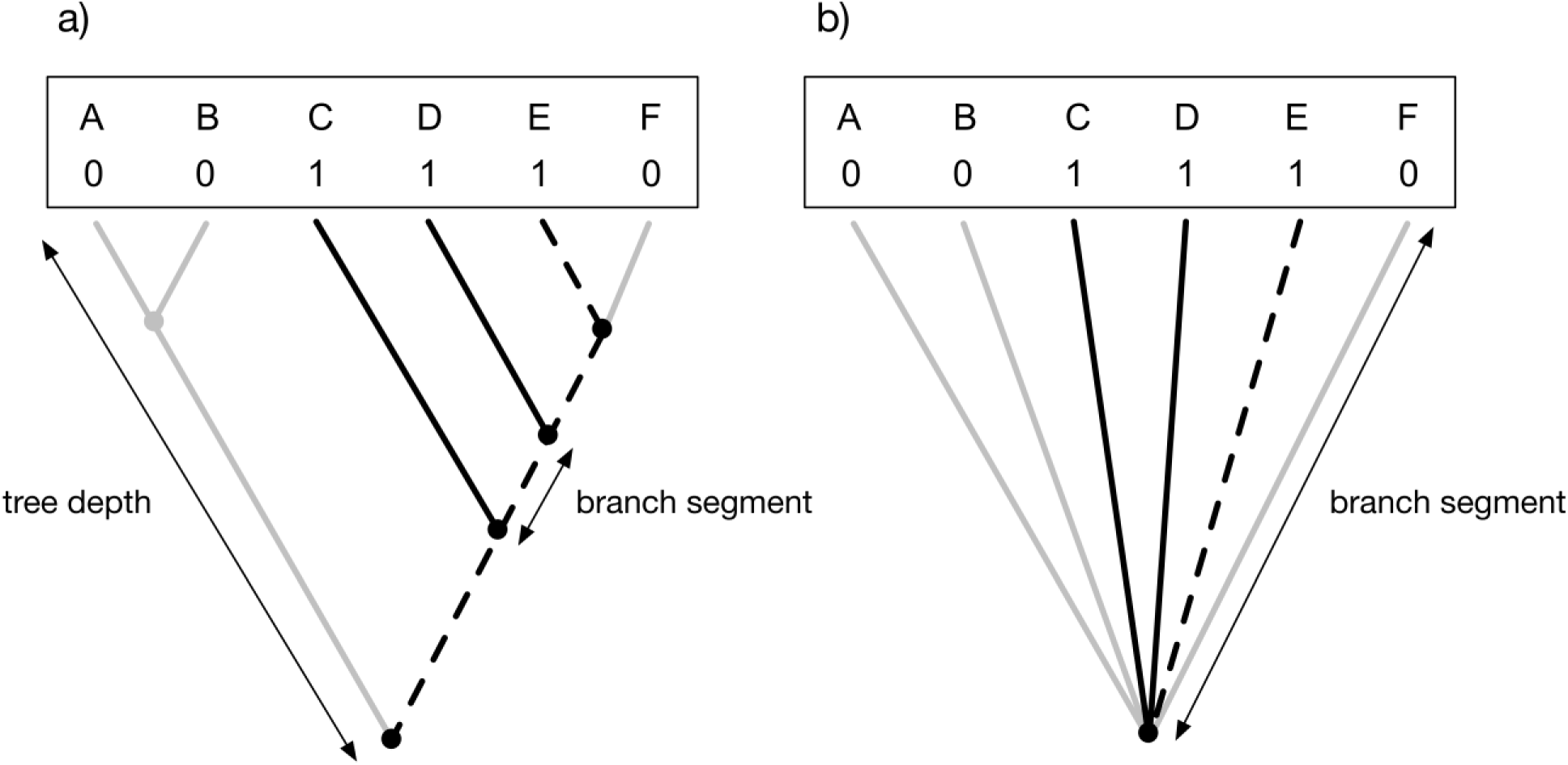
Tree-based diversity where diversity is defined as the total path length connecting a set of species to the root of the tree. In this hypothetical example, the presence or absence of species A-F in a site are shown by 1 or 0, respectively. Internal nodes are represented as circles and branch segments are lines that connect to tips to nodes or nodes to nodes. The tree-based diversity of the site is the sum of the lengths of the branch segments that connect the species present in the site to each other and to the root of the tree (total path length shown in black). The particular set of branch segments that connect species E to the root of the tree are shown as dashed lines. In tree a), species differ in their similarity to each other and thus share, or don’t share, particular branch segments. Tree b) shows how species richness can be viewed as a special case of tree-based diversity where species do not share branch segments and where tree depth is equal to 1.

For tree-based diversity, the expected complementarity (*E*[*ΔD*(*i*, *m*)]) of site *i* is the sum of the expected *contribution* of each branch segment (*k*) present in site *i* to the complementarity of site *i* (Faith et al. 2004). A branch segment is present in a site if it contributes to the tree-based diversity of a site - that is, it constitutes some portion of the path connecting a species present in the site to the root of the tree (Figure 1). The expected contribution of branch segment *k* is the product of its length (*l*_*k*_) and the probability (*q*_*ik*_) that it is absent from a subset of sites (*M* where *i* ∉ *M*) of size *m*, drawn at random and without replacement from *N* (equation 3).

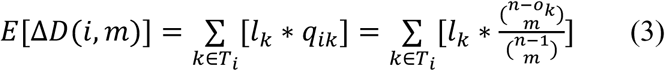

Where *o*_*k*_ is the frequency (no. of sites) with which branch segment *k* occurs in sites across the set, *N*, and *T*_*i*_ is the subset of branch segments in a tree (*T* where *T*_*i*_ ⊂ *T*) that connects the species of site *i* to each other and to the root (Figure 1). As with equation 2, equation 3 has a direct relation to rarefaction and specifically to rarefaction of Phylogenetic Diversity (Chao et al. 2015; Nipperess & Matsen 2013; Nipperess 2016; Tsirogiannis et al. 2012). Note that when the tree-based diversity of a site is represented by a “star phylogeny” where no species share branch segments and all branch segments have a length of 1 (Figure 1), then *n*_*j*_ = *o*_*k*_ and *l*_*k*_ = 1, and equation 3 is identical to equation 2.

The frequency (*o*_*k*_) with which a branch segment (*k*) occurs across a set of sites (*N*) is given by equation 4.

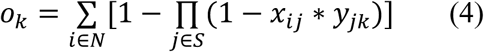

Where *y*_*jk*_ is a binary value that equals 1 if branch segment *k* constitutes some portion of the path connecting species *j* to the root of the tree, *T*, and is zero (0) otherwise.

To explore the full range of possible complementarity values for some site, we can plot expected complementarity against the number of randomly chosen sites (*m*) as a “complementarity curve”. Figure 2 shows an example for a dataset of land birds of the Sipoo archipelago, Finland (Simberloff & Martin 1991). The complementarity curve visualises the conservation importance of a site for all possible combinations from *m* = 0 to *m* = *n* – 1. When *m* = 0, expected complementarity is equal to the diversity of the site because all species present in the site contribute to overall diversity. When *m* = *n* – 1, expected complementarity is equal to the endemism of the site because only species that are uniquely found in the site will make any contribution to overall diversity.

**Figure 2:**
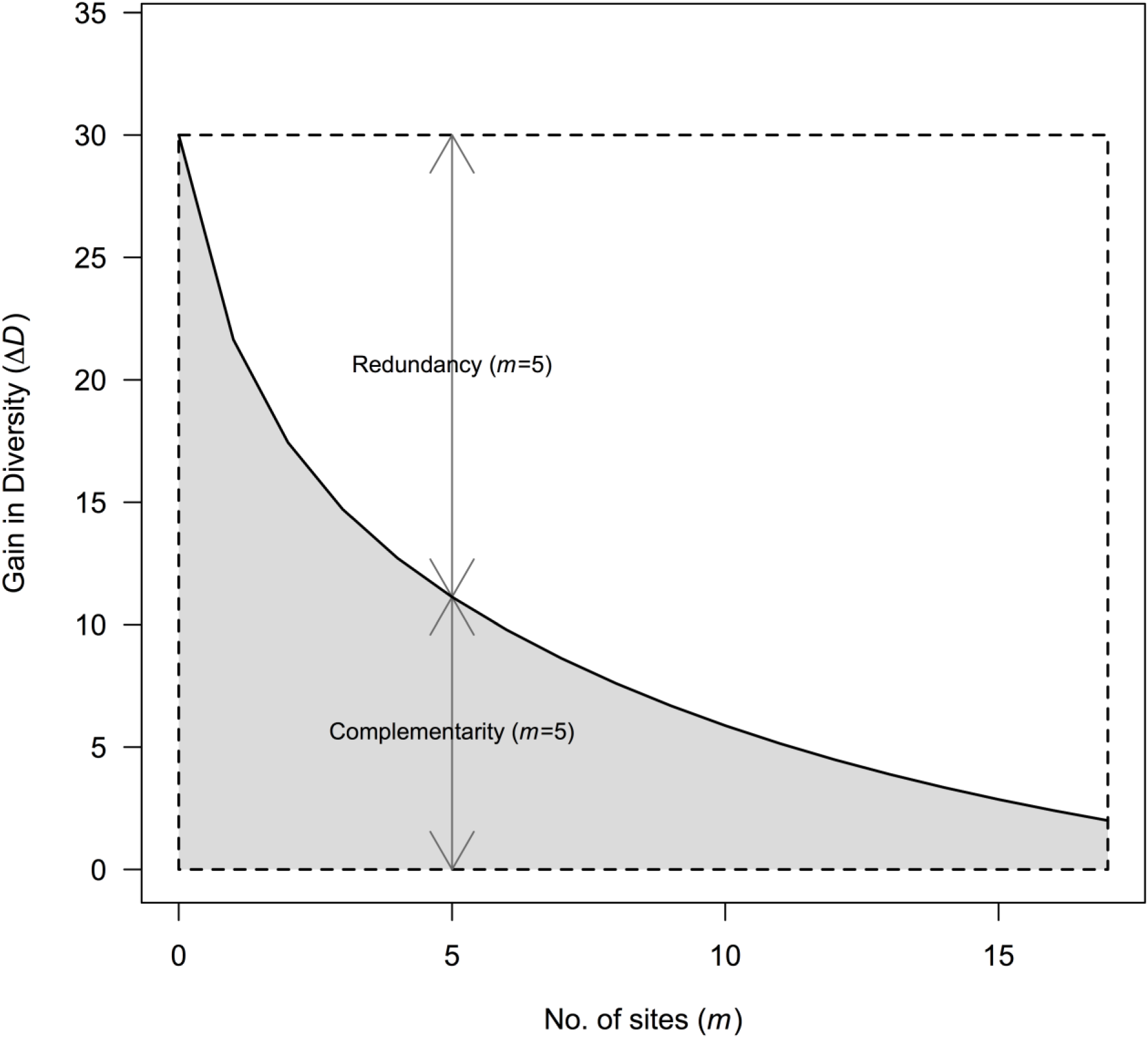
Complementarity curve for the land bird assemblage of the island of Kaunissri in the Sipoo archipelago, southern Finland. The curve shows expected complementarity in species richness of Kaunissri in comparison to all possible subsets (of size, *m*) of 17 other islands in the archipelago. Total species richness of the island (30 spp.) is the leftmost point of the curve and the number of species unique to the island (2 spp.) is the rightmost point of the curve. The expected irreplaceability of Kaunissri’s bird assemblage is the area under the curve (shaded grey), while the remaining area in the dashed box is the expected replaceability. Complementarity and corresponding redundancy for a particular point on the curve (*m*=5) is shown. Data sourced from Simberloff & Martin (1991).

If we interpret the irreplaceability of a site as its overall contribution to diversity across the full range of conservation scenarios (all possible combinations of sites), then it can be defined as the area under its complementarity curve (Figure 2). Because the values for *m* are discrete, irreplaceability (*AUC*[*ΔD*(*i*)]) can be simply calculated by dividing the area under the curve into polygons and summing their area (equation 5).

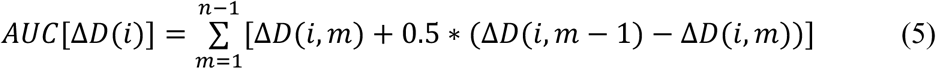

Finally, we can define two further properties for a specific site, *Redundancy* and *Replaceability*, which are simply the mathematical complements of the corresponding values for complementarity and irreplaceability, respectively (Figure 2). Redundancy is the extent to which species present in site *i* are represented in other sites in a set of *m* sites and is simply the difference between the diversity (*D*(*i*)) of the site and its complementarity (*ΔD*(*i*, *m*)) for some value of *m* (where *m* > 0). Replaceability is the overall redundancy of site *i* for all values of *m* (i.e. the area *above* the complementarity curve) and can also be determined by simple subtraction (Equation 6).

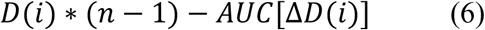

## Discussion

Generalised Complementarity Analysis (GCA) provides a probabilistic framework for understanding the importance of any site to the conservation of biodiversity across some spatial domain. The core concepts of complementarity and irreplaceability are given explicit mathematical definitions that are consistent with prior usage. Essentially, complementarity becomes the expected gain in diversity of adding a focal site to an existing set of sites and irreplaceability becomes the overall expected gain for all possible sets of sites in the domain.

As defined by GCA, complementarity and irreplaceability essentially measure the endemism of a site. In fact, in the case where a focal site is added to all other sites in the domain (i.e. the end of the complementarity curve where *m* = *n* – 1), expected complementarity is exactly equal to absolute endemism (no. of species found only in the site). For the rest of the curve, complementarity can be considered a kind of weighted endemism (Rosauer et al. 2009; Crisp et al. 2001). Rather than count only species that occur uniquely in a site (as in absolute endemism), each species (or other unit of biodiversity) is assigned a weight that is the inverse of that species' range (area or count of sites occupied). Thus, species with wide ranges are assigned a small weight and species with small ranges are assigned a large weight. Weighted endemism is then the sum of these weights for species present at a site. The benefit function of the Zonation algorithm (Moilanen et al. 2005) uses a similar weighting for species' ranges. Expected complementarity in GCA is also a sum of species’ weights, where the weight is defined as the probability that the species will be found uniquely in the focal site. The more sites in which some species is found, the less likely it will be absent from a randomly chosen subset and the less likely it will be uniquely found in the focal site. Thus, expected complementarity can be interpreted as a probabilistic version of weighted endemism and will scale monotonically with other endemism scores.

Generalised Complementarity Analysis is superior to previous conservation planning approaches in that is is an exact analytical solution. This means that expected complementarity is known exactly (i.e. for *all* possible combinations of *m* sites, rather than a sample) and the value can be quickly calculated (directly from the species’ probabilities). The alternative method of deriving an expected complementarity would be to either randomly resample a large number of times (Williams et al. 2006) or to iteratively search through possible site combinations for optimal combinations (Moilanen et al. 2005; Rebelo & Siegfried 1992; Ball et al. 2009). GCA has the advantage of being considerably more computationally efficient than these approaches and is thus scalable to very large numbers of sites. This is especially important when trying to develop gridded maps of conservation priority. Even for a tiny 5 x 5 grid of 25 sites, there are literally millions of possible combinations against which any one site could be compared. Given the growing emphasis on global conservation prioritisations (Montesino Pouzols et al. 2014; Pollock et al. 2017), GCA provides a rapid means of producing gridded maps that can be simultaneously high resolution and of large extent.

However, GCA is not intended as a replacement for heuristic conservation planning algorithms but rather as a useful supplementary tool. GCA does not find optimal combinations of sites for conservation, nor does it account for the differential costs of acquiring sites, nor does it assign weights to sites or combinations of sites based on their capacity to produce connected reserve networks. Existing algorithmic solutions do account for these things and more. Instead, GCA is a useful null model that identifies the overall importance of a site, across all possible scenarios, rather than its contribution to a specific set of sites selected by an algorithm. This allows planners to determine to what extent a site is *generally* important versus *specifically* important and gives an insight into where optimised algorithmic solutions are placed within a wider universe of possible solutions. Because GCA is an analytical solution, insight can be gained without having to exhaustively sample that wider solution space. Further, any site that is generally important will always be specifically important, regardless of conservation plan, because it will, by definition, contain geographically rare (and possibly functionally and evolutionarily distinct) species. Identifying such generally important sites is thus of value in itself, even in the absence of more detailed analyses providing heuristic optimised solutions.

By identifying truly irreplaceable sites, regardless of conservation scenario, GCA opens up new areas of conservation research. For example, conservation planning needs comprehensive data on species distribution and so is typically only conducted on well-known taxa such as mammals or birds. This is concerning because such taxa constitute a very small proportion of biodiversity and yet we assume, without sufficient evidence, that conserving areas of significance for well-known groups will be adequate for protecting poorly known groups, such as insects (Nipperess et al. 2012). Our confidence in this assumption can be assessed if we are able to compare maps of irreplaceability for multiple taxa (Williams et al. 2006) and, more significantly, develop statistical models that predict irreplaceability based on environmental and other spatial variables (Ferrier et al. 2004). Despite the considerable gaps in our understanding of the distribution of biodiversity, the need to develop effective conservation plans is urgent and the tools for doing so are increasingly at hand.

## Acknowledgments

This manuscript was conceived and written without any grant funding, salary or other financial support.

## References

Asmyhr, M.G. et al., 2014. Systematic Conservation Planning for Groundwater Ecosystems Using Phylogenetic Diversity. PLoS One, 9(12), e115132.

Ball, I.R., Possingham, H.P. & Watts, M., 2009. Marxan and relatives: Software for spatial conservation prioritisation. Spatial conservation prioritisation Quantitative methods and computational tools, pp. 185–195.

Cardoso, P. et al., 2014. Partitioning taxon, phylogenetic and functional beta diversity into replacement and richness difference components. Journal of Biogeography, 41(4), pp. 749–761.

Chao, A. et al., 2015. Rarefaction and extrapolation of phylogenetic diversity. Methods in Ecology and Evolution, 6(4), pp. 380–388.

Crisp, M.D. et al., 2001. Endemism in the Australian flora. Journal of Biogeography, 28(2), pp. 183–198.

Faith, D.P., 1992. Conservation evaluation and phylogenetic diversity. Biological Conservation, 61, pp. 1–10.

Faith, D.P., Reid, C.A.M. & Hunter, J., 2004. Integrating phylogenetic diversity, complementarity, and endemism for conservation assessment. Conservation Biology, 18(1), pp. 255–261.

Ferrier, S. et al., 2004. Mapping more of terrestrial biodiversity for global conservation assessment. BioScience, 54(12), pp. 1101–1109.

Hurlbert, S., 1971. The nonconcept of species diversity: a critique and alternative parameters. Ecology, 52(4), pp. 577–586.

Lande, R., 1996. Statistics and partitioning of species diversity, and similarity among multiple communities. Oikos, 76(1), pp.5–13.

Margules, C. & Pressey, R., 2000. Systematic conservation planning. Nature, 405, pp. 243–253.

Moilanen, A. et al., 2005. Prioritizing multiple-use landscapes for conservation: methods for large multi-species planning problems. Proceedings of the Royal Society, Biological sciences, 272(1575), pp. 1885–1891.

Montesino Pouzols, F. et al., 2014. Global protected area expansion is compromised by projected land-use and parochialism. Nature, 516(7531), pp. 383–386.

Nipperess, D.A. et al., 2012. Plant phylogeny as a surrogate for turnover in beetle assemblages. Biodiversity and Conservation, 21(2), pp. 323–342.

Nipperess, D.A., 2016. The Rarefaction of Phylogenetic Diversity: Formulation, Extension and Application. Biodiversity Conservation and Phylogenetic Systematics, 14, pp. 197–217.

Nipperess, D.A. & Matsen, F.A., 2013. The mean and variance of phylogenetic diversity under rarefaction. Methods in Ecology and Evolution, 4, pp. 566–572.

Olszewski, T.D., 2004. A unified mathematical framework for the measurement of richness and evenness within and among multiple communities. Oikos, 104(2), pp. 377–387.

Petchey, O.L. & Gaston, K.J., 2002. Functional diversity(FD), species richness and community composition. Ecology Letters, 5, pp. 402–411.

Pollock, L.J. et al., 2015. Phylogenetic diversity meets conservation policy: small areas are key to preserving eucalypt lineages. Philosophical Transactions of the Royal Society of London B, 370(1662), 20140007.

Pollock, L.J., Thuiller, W. & Jetz, W., 2017. Large conservation gains possible for global biodiversity facets. Nature, 546(7656), pp. 141–144.

Pressey, R.L., Johnson, I.R. & Wilson, P.D., 1994. Shades of irreplaceability: towards a measure of the contribution of sites to a reservation goal. Biodiversity and Conservation, 3(3), pp. 242–262.

Rebelo, A.G. & Siegfried, W.R., 1992. Where should nature reserves be located in the Cape Floristic Region, South Africa? Models for the spatial configuration of a reserve network aimed at maximizing the protection of floral diversity. Conservation Biology, 6(2), pp. 243–252.

Rosauer, D. et al., 2009. Phylogenetic endemism: a new approach for identifying geographical concentrations of evolutionary history. Molecular Ecology, 18(19), pp. 4061–4072.

Rosauer, D.F. & Mooers, A.O., 2013. Nurturing the use of evolutionary diversity in nature conservation. Trends in Ecology & Evolution, 28(6), pp. 322–323.

Simberloff, D. & Martin, J.L., 1991. Nestedness of insular avifaunas: simple summary statistics masking complex species patterns. Ornis Fennica, 68, pp. 178–192.

Tsirogiannis, C., Sandel, B. & Cheliotis, D., 2012. Efficient Computation of Popular Phylogenetic Tree Measures. Algorithms in Bioinformatics, 7534, pp. 30–43.

Williams, P. et al., 2006. Complementarity analysis: Mapping the performance of surrogates for biodiversity. Biological Conservation, 128, pp. 253–264.

